# Transcriptomic sexual conflict at two evolutionary timescales revealed by experimental evolution in *Caenorhabditis elegans*

**DOI:** 10.1101/2023.08.09.552689

**Authors:** Katja R. Kasimatis, John H. Willis, Christine A. Sedore, Patrick C. Phillips

**Affiliations:** Institute of Ecology and Evolution, University of Oregon, Eugene, Oregon, United States of America; Department of Biology, University of Virginia, Charlottesville, Virginia, United States of America

## Abstract

Sex-specific regulation of gene expression is the most plausible way for generating sexually differentiated phenotypes from an essentially shared genome. However, since genetic material is shared, sex-specific selection in one sex can have an indirect response in the other sex. From a gene expression perspective, this tethered response can move one sex away from their wildtype expression state and impact potentially many gene regulatory networks. Here, using experimental evolution in the model nematode *Caenorhabditis elegans,* we explore the coupling of direct sexual selection on males with the transcriptomic response in males and females over microevolutionary timescales to uncover the extent to which post-insemination reproductive traits share a genetic basis between the sexes. We find that differential gene expression evolved in a sex-specific manner in males, while in females indirect selection causes an evolved response. Almost all differentially expressed genes were downregulated in both evolved males and females. Moreover, 97% of significantly differentially expressed genes in males and 69% of significantly differentially expressed genes in females have wildtype female-biased expression profile. Changes in gene expression profiles were likely driven through *trans*-acting pathways that are shared between the sexes. We found no evidence that the core dosage compensation machinery was impacted by experimental evolution. Together these data suggest a de-feminization of the male transcriptome and masculinization of the female transcriptome driven by direct selection on male sperm competitive ability. Our results indicate that on short evolutionary timescales sexual selection can generate putative sexual conflict in expression space.

## INTRODUCTION

Sexual dimorphism, ubiquitous across multicellular organisms, is rooted in the seeming conundrum that the sexes can be dramatically different yet share largely the same genome [1,2]. On a functional level, this means that gene expression must be tightly regulated in a sex-specific manner [3,4]. On an evolutionary scale, it also means that selection acting differentially on one sex can have an immediate correlated response to selection on attributes of the other sex [5,6].

This correlation underpins the classic Fisherian model of runaway sexual selection, as direct selection acting on a male display trait generates self-reinforcing indirect selection on female preference if they share some form of joint inheritance, either through pleiotropy or linkage [7–9]. Indirect selection can in principle generate a correlated evolutionary response on any number of female traits, however, particularly on polygenic traits for which alleles are more likely to be pleiotropic [10]. For example, males of *Pteridophora alberti*, the King of Saxony bird-of-paradise, have highly elongated feathers equal to several body lengths on either side of their heads. Curiously, females display similar, but much smaller, feathers on their heads, possibly as residual responses to sexual selection in males [11,12]. Much like these birds, opposite-sex perturbation would be expected to be fairly short-lived on an evolutionary timescale, as natural selection against misexpression in the wrong sex leads to heightened sexual dimorphism [13].

The degree to which mating traits have additive genetic covariance, then, determines the opportunity for sexual conflict [14,15]. While the evolution of sexual dimorphism has been extensively studied for macro-level secondary sexual characteristics, such as plumage, territorial behavior, and body size, much less is known at the molecular level, particularly about the coupling between the primary domains of dimorphism: the response to sexual selection at the evolutionary level and the evolution sex-specific gene regulation at the functional level. We address that coupling here with an experimental evolution framework using the nematode *Caenorhabditis elegans*.

In internally fertilizing species, there is an additional layer to the reproductive process to those classically studied under sexual selection, namely post-insemination reproductive interactions. Here the male “trait” of interest can be any cell or protein that affects male fertilization success, while female “preference” is realized in any tissue, cell, or protein that interacts with the male ejaculate to affect female fertilization success. Sexual selection on male reproductive proteins has been inferred through molecular evolution analyses [16–19]. These studies indicate that many seminal fluid proteins evolve rapidly due to sexually antagonistic coevolution, which in turn implies that direct selection is acting on both female and male post-insemination traits. However, identifying the specific female traits with which these male reproductive proteins interact has proved challenging. In this case, transcriptomic analysis can be useful for identifying wide-spread changes in gene expression after mating to uncover polygenic responses to selection within and across generations.

Few studies have measured direct selection on male post-insemination traits and the correlated response in females in a similar manner to male display and female preference traits. We previously used experimental evolution to select for increased sperm competitive ability in *C. elegans* males [20]. Experimental evolution provides an opportunity to reproducibly isolate directional selection on male characteristics from the potentially correlated responses in females using timescales when the opportunity for sexual conflict is likely to be highest. We found that after 30 generations of evolution, males showed a strong, rapid response to post-insemination selection at both the phenotypic and genomic levels [20]. Specifically, post-insemination reproductive success increased by greater than four-fold in the evolved populations. This reproductive fitness increase was underlain by a polygenic response of 82 significance peaks represented by 57 genes, 10 intergenic regions, and 15 putative pseudogenes. Together we found that when the androdioecous *C. elegans* mating system is genetically engineered to reflect the ancestral diecious mating state, there is overwhelming selection to improve maleness.

Here we test if this strong, direct selection acting on male sperm competitive ability had an indirect impact on females. Specifically, we quantify the impact of direct post-insemination sexual selection on the transcriptomes of mated ancestral and evolved males and compare how selection impacts the transcriptome relative to the genome. Additionally, we examine the transcriptomes of mated females using a fully factorial crossing design of ancestral and evolved individuals to determine: i) whether strong, direct selection on sperm competition generates a response in females, and if so, ii) is the female response evolved or plastic, and iii) is the female response indicative of sexual conflict.

## RESULTS

### A strong-inference framework for expression evolution

We selected four experimental evolution replicates that evolved under direct selection on sperm competitive ability for 30 generations to test for changes in the transcriptional profile of mated males and mated females. Ancestral and evolved males were crossed with the corresponding generation of ancestral or evolved females to assess how direct selection on males altered the male transcriptome. For females, we used a full factorial crossing design of ancestral and evolved females mated with ancestral and evolved males to determine if the female transcriptome changes and whether those changes were a result of selection (direct or indirect) or female plasticity. Populations of individuals for each cross were mated for 48 hours after which individuals of the focal sex were isolated for transcriptomic analysis. The ancestor_F_-ancestor_M_ cross represents the baseline transcriptomic profile of both males and females after mating before experimental evolution occurred. The evolved_F_-evolved_M_ cross represents the transcriptomic profile of males and females after 30 generations of evolution. If direct sexual selection on males had no impact on a gene’s expression level, then there would not be differential expression between the ancestor_F_-ancestor_M_ and evolved_F_-evolved_M_ cross for that gene. In males, such genes would not contribute to the phenotype under selection, namely sperm competitive ability. In females, such genes would have no genetic covariance between females and males.

If instead, in females, a gene is differentially expressed between the ancestor_F_-ancestor_M_ and evolved_F_-evolved_M_ crosses, then females are responding to the sexual selection exerted on males. If the female transcriptional changes are a plastic response to increased sperm competitiveness, then male generation should be a major predictor, that is females of the evolved_F_-ancestor_M_ cross should have expression profiles similar to the ancestor_F_-ancestor_M_ cross and females of the ancestor_F_-evolved_M_ should have expression profiles similar to the evolved_F_-evolved_M_ cross. Alternatively, if female transcriptional changes are a result of either direct selection in females induce by sperm competitiveness or indirect selection generated by genetic covariation between the sexes, then the expectation is that ancestral females will have similar expression profiles and evolved females will have similar expression profiles, regardless of male generation.

### Direct selection on males repeatably downregulates the evolved male transcriptome

We analyzed changes in male gene expression for each experimental evolution replicate separately (Fig S1, Fig S2). Principal component analysis (PCA) was performed to compare samples basted on male generation (*i.e.*, G0 ancestor or G31 evolved). In each replicate, the first principal component partitioned samples by male generation and explained 68% to 82% of the total variance in gene expression (Fig S1). Thus, there was a strong effect of direct selection on the male transcriptome. Differential gene expression analysis showed an overall signature of downregulation with 50% (Replicate B2) to 91% (Replicate A3) of genes being downregulated in evolved males. However, the degree of overlap between significantly differentially expressed genes (DEs) varied greatly among replicates (Fig S3A). Ninety-eight percent of DEs in Replicate A3 were significant in at least one other replicate, while only 60% of DEs in Replicate A2 were shared with at least one other replicate. Replicate A2 also had the most DEs (n = 2,181) and showed the strongest increase in post-mating reproductive success [20], indicating replicate-specific variation in the transcriptional response.

Differential evolution between experimental evolution replicates is not surprising and could be cause by multiple processes, including selection acting on different segregating genetic variants and genetic drift. To focus on the consistent and repeatable gene expression changes that are most likely linked with the phenotype under selection, we combined all experimental evolution replicates. The final combined dataset in males consisted of three independent ancestor_F_-ancestor_M_ crosses, and 12 independent evolved_F_-evolved_M_ crosses.

PCA again clustered samples by male generation, such that all ancestral males clustered and all evolved males clustered (Fig 1A). The first principal component explained 42% of the variation in total gene expression and separated males by generation. The combined analysis, therefore, supports the individual replicate analysis and shows a strong effect of direct selection on the male transcriptome. We analyzed differential gene expression using a generalized linear model framework using male generation and correcting for average differences among evolutionary replicates. Differential expression analysis showed 589 genes to be significantly differentially expressed based on male generation of which 125 had at least a two-fold change in expression (Fig 1B; File S1). All but 18 DEs were represented in individual replicate analyses, which indicates that the transcriptional changes identified are likely due to selection (Fig S3B). Replicate did not contribute to significant differential gene expression (File S1).

**Figure 1.**
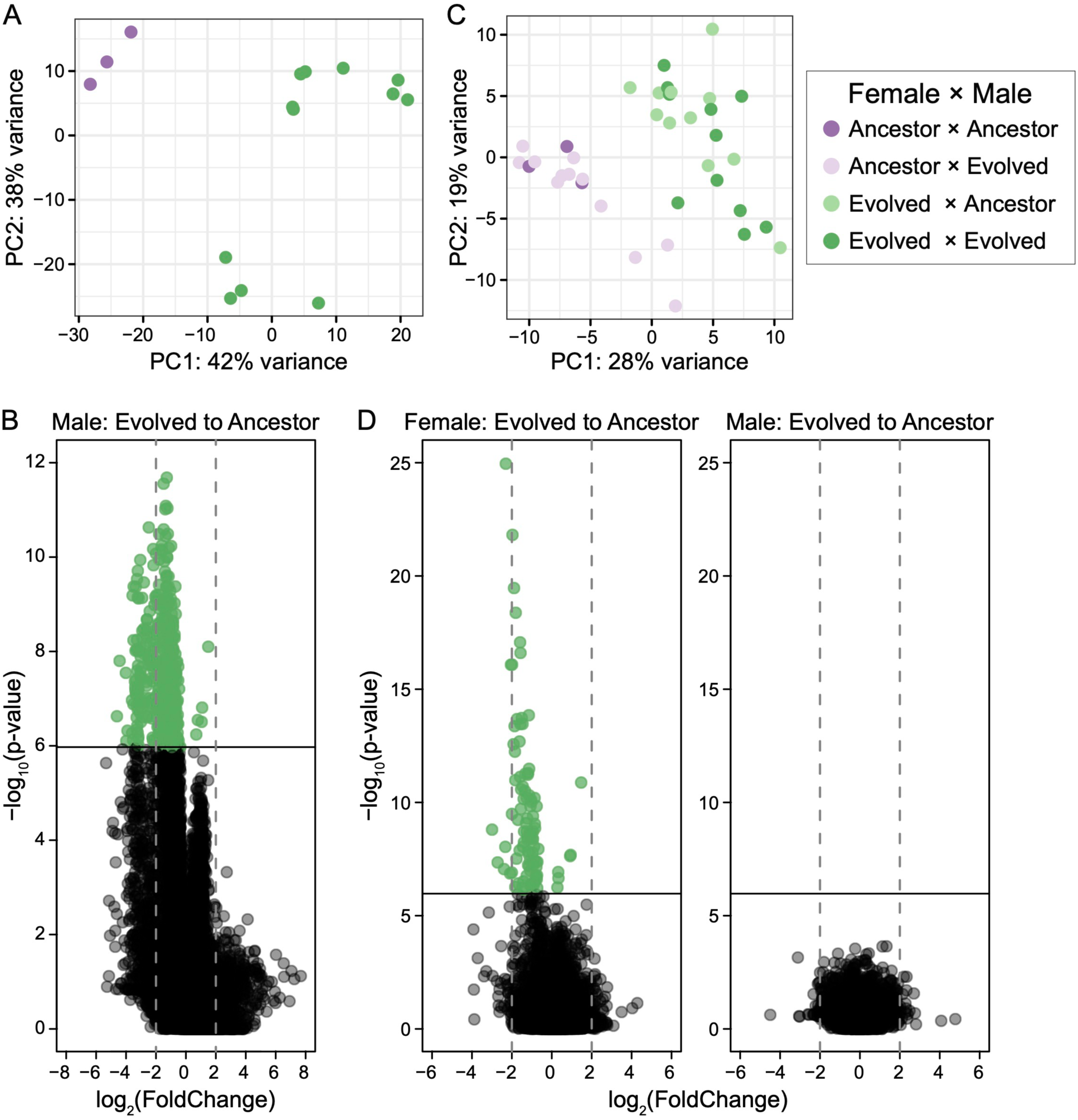
Differential gene expression is partitioned by ancestral (purple) and evolved (green) individuals. **A)** Principal component analysis of the variance stabilized dataset partitions male generation along PC1. **B)** Differential gene expression was analyzed based on male generation and experimental evolution replicate. Only the male term contributed to gene expression changes (File S1). Genes with a significant change in expression from ancestral to evolved males (shown in green) were determined using a Bonferroni cut-off. Positive expression changes correspond to upregulation in evolved males, while negative expression changes correspond to downregulation in evolved males. **C)** Principal component analysis of the variance stabilized dataset partitions female generation along PC1. **D)** Differential gene expression was analyzed based on female generation, male generation, and experimental evolution replicate. Only the female term contributed to gene expression changes. Genes with a significant change in expression from ancestral to evolved females (shown in green) were determined using a Bonferroni cut-off. Positive expression changes correspond to upregulation in evolved females, while negative expression changes correspond to downregulation in evolved females.

Only five genes were significantly upregulated (mean fold increase = 1.0) in evolved males relative to ancestral males. They are located on autosomes and do not have ontology terms that clearly connect them to reproduction. The other 584 genes (99%) were downregulated (mean fold decrease = 1.5) in evolved males relative to ancestral males. The widespread signature of downregulation is not an artifact of biased count distribution (Fig S4A). Rather, the genes which are highly upregulated in evolved males tended to occur in specific experimental evolution replicates, highlighting the importance of multiple levels of replication when identifying consistent patterns of selection. To determine if SNPs in the coding region or the *cis*-regulatory region of these genes were responsible for the changes in expression, we examined allele frequencies in our previously published genomic mapping data for these populations [20]. None of the DEs had significant allele frequency changes in the coding region. Two genes (*apn-1* and *apa-2*) had SNPs in the *cis*-coding region, however, none of these SNPs showed a significant allele frequency change over the course of experimental evolution (File S2). Together, these data indicate that direct selection on males is downregulating the male transcriptome through *trans*-acting pathways.

To assess the functional pathways in which these downregulated genes were involved, we examined their associated gene ontology (GO) terms and KEGG Pathways. GO enrichment analysis identified 20 ontology terms that were significantly enriched, many of which were related to some form of binding: DNA binding (n = 4), RNA binding (n = 2), and protein binding (n = 1) (File S3). Eight KEGG pathways were significantly enriched, including four in the replication and repair class, three in the metabolism class, and one in the endocrine system class (File 3).

### Female gene expression changes correlate with males based on female generation

First, to assess variation in the evolutionary response, we analyzed changes in female expression for each experimental evolution replicate separately. PCA clustered females by generation in each replicate (Fig S5). The first principal component explained 30% to 46% of the total variance in gene expressed based on female generation alone. Male generation did not contribute to variance partitioning in the first two principal components. Replicated B3 had two outlier transcriptome replicates, which were censored in the downstream analyses (Fig S5). Differential expression analysis supported that female generation alone determined expression changes as the male term did not identify DEs (Fig S6). Female DEs recapitulated the male signature of downregulation with 92% to 96% of DEs being downregulated in evolved females. The majority of DEs overlapped with at least one other experimental evolution replicate, suggesting less replicate-specific evolution in females than in males (Fig S3C).

Although the separate analysis of each replicate helps to highlight variance in evolution response, to more formally address evolutionary changes in the transcriptome that are repeatable across replicates, we used a combined analysis after correcting for among-replicate variance. The final censored, combined dataset in females therefore consisted of three independent ancestor_F_-ancestor_M_ crosses, 12 independent ancestor_F_-evolved_M_ crosses, 11 independent evolved_F_-ancestor_M_ crosses, and 11 independent evolved_F_-evolved_M_ crosses. PCA of the full dataset again clustered samples by female generation along the first principal component, which explained 28% of the total variance in gene expression levels (Fig 1B). Male generation did not contribute to variance partitioning in the first two principal components, which captured 46% of the total variance. Thus, the combined PCA supports the individual replicate analyses and indicates that the female response is evolved rather than a plastic response based on male generation.

To formally examine potential differences of ancestral and evolved males on the female transcriptional response, as well as the influence of evolutionary response to heightened sperm competition of the females themselves, differential expression for individual genes was analyzed using a generalized linear model using female and male generation, correcting for average differences among evolutionary replicates. Neither male generation nor replicate contributed to significant changes in gene expression (Fig 1D; File S1). Rather, female generation determined differential gene expression, again demonstrating an evolved expression response (Fig 1D). One hundred genes showed significant differential gene expression of which eight genes had at least a two-fold change in expression (Fig 1D). Only six genes were significantly upregulated in evolved (mean fold increase = 0.7) females relative to ancestral females. Five of these genes are uncharacterized in function. The other 94 DEs were downregulated (mean fold decrease = 1.3) in evolved females relative to ancestral females (Fig 1D). Again, widespread downregulation is not an artifact of biased count distribution (Fig S4B). None of the significantly differentially expressed genes had significant allele frequency changes in the coding region or the *cis*-regulatory region (File S1). Thus, changes in gene expression were again likely regulated through *trans*-acting pathways. No GO terms or KEGG pathways were significantly enriched in female DEs.

### Indirect selection on females drives transcriptome evolution

Since the female transcriptome response is evolved, we next examined whether the changes in female expression were due to direct selection in response to increased sperm competitiveness or indirect selection as a correlated response to the direct selection acting on males. To this end, we examined at the correlation in differential gene expression changes between the sexes. Genes that were only significantly differentially expressed in males (*i.e.*, unique male DEs) were not correlated with female changes in gene expressed (n = 565, p = 0.27, adjusted R^2^ = 0.0004; Fig 2). Rather, these genes changed expression in males alone. This widespread male-specific transcriptomic response indicates that direct selection on males predominantly changed the male transcriptome in a sex-specific manner.

**Figure 2.**
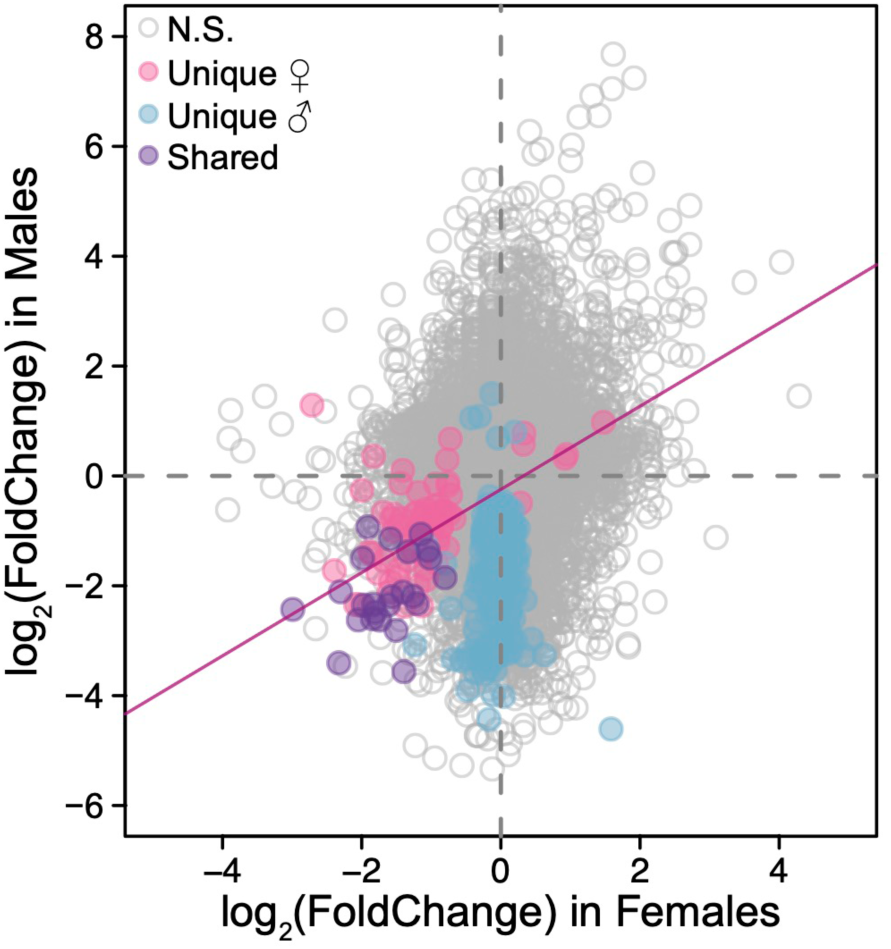
A comparison of changes in gene expression levels between females and males. Positive expression changes correspond to upregulation in evolved females/males, while negative expression changes correspond to downregulation in evolved females/males. Genes that are not significantly differentially expressed in either sex are shown in gray. Genes that are significantly differentially expressed in males but not in females (*i.e.*, unique male) are shown in blue. Genes that are significantly differentially expressed in females but not in males (*i.e.*, unique female) are shown in pink. Genes that are significantly differentially expressed in both sexes (*i.e.*, shared) are shown in purple. There is a significant relationship between the change in gene expression in females and males for all female DEs (F_1,98_ = 42.16, p < 0.001, adjusted R^2^ = 0.29).

Genes that were only significantly differentially expressed in females (*i.e.*, unique female DEs) were, however, significantly correlated with corresponding changes in male expression (n= 76, F_4,74_ = 21.3, p < 0.001, adjusted R^2^ = 0.21). Therefore, these genes are largely being downregulated in females and males, though not significantly so in males (Fig 2). This correlation holds when all female DEs – unique and shared between the sexes (n = 24) – are included (F_1,98_ = 42.16, p < 0.001, adjusted R^2^ = 0.29). Together, these data support that indirect selection on females was driving changes in the female transcriptome as female expression changes were not independent from those in male, while male expression changes occurred in a sex-specific manner.

### Correlated selection de-feminizes the transcriptome of males and females

We hypothesized that selection was driving this widespread downregulation via either female-biased or non-sex biased genes. In either case, the transcriptomic effect would be a less female-like and a more male-like transcriptome to support the increase in male reproductive fitness. To test this hypothesis, we examined the relationship between changes in gene expression after experimental evolution and sex-biased expression in wildtype females and males [21]. The majority (n = 571) of male DEs showed some degree of female-biased expression in wildtype individuals (Fig 3A). A strong negative correlation exists between wildtype sex-bias and gene expression changes in males, such that more female-biased genes in wildtype were more strongly downregulated after experimental evolution regardless of whether the gene was located on an autosome or the X chromosome (F_1,568_ = 1562, p < 0.0001, adjusted R^2^ = 0.73). The reduction in female-bias indicates a de-feminization of the male transcriptome.

**Figure 3.**
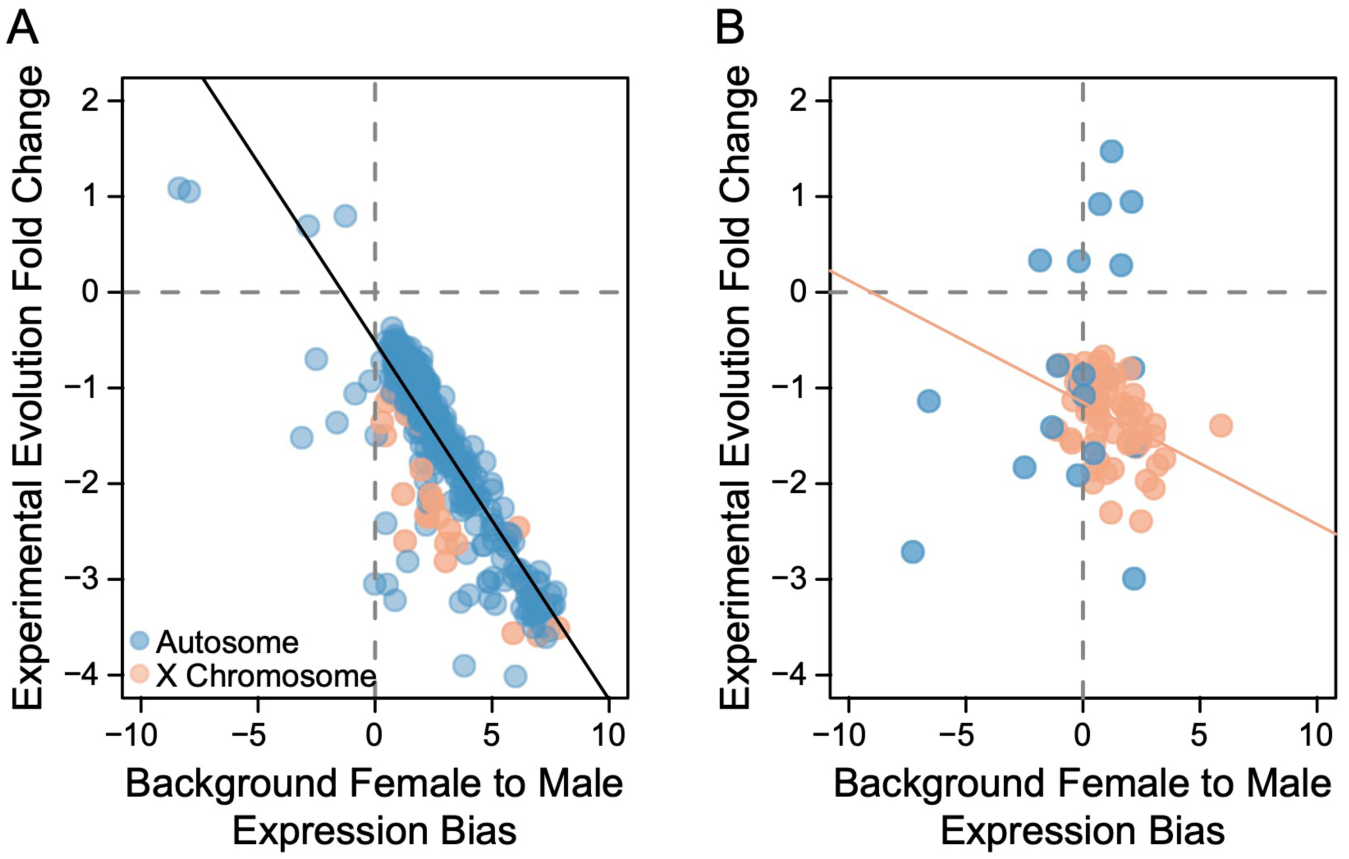
Downregulated genes are female-biased. **A)** There is a significant negative correlation between wildtype sex-bias in expression and changes in expression after experimental evolution in males. This relationship holds regardless of genomic location. Positive wildtype gene expression values indicate female-biased expression and negative values indicate male-biased expression. Both sex-biased expression (female to male) and experimental evolution fold change are plotted on a log_2_ scale. **B)** There is a negative correlation between significant differentially expressed genes located on the X chromosome (shown in coral) and sex-biased expression in females. Only 22 genes were located on autosomes (shown in blue).

Similarly, the majority (n = 69) of female DEs also showed some degree of female-biased expression in wildtype individuals (Fig 3B). Interestingly, 67 of the downregulated genes (81%) were located on the X chromosome, of which 57 were female-biased (Fig 3B). There is a negative relationship between sex-biased expression and differential gene expression on the X chromosome, such that more female-biased genes had a greater fold decrease in expression (F_1,65_ = 12.4, p < 0.001, adjusted R^2^ = 0.15). This relationship suggests a pleiotropic effect in a *trans*-regulatory pathway impacting regulation of X-linked genes. No relationship exists between autosomal genes and differential gene expression (p = 0.14), though the low number of autosomal genes likely makes this comparison under-powered. Together, these data support a correlated expression response in females and indicate a skew towards a less female-like transcriptome in evolved females.

Significantly more DEs are located on the X chromosome in females than in males (χ^2^ = 319.5, d.f. = 1, p < 0.0001) (Fig S7). One major *trans*-regulatory pathway that could explain the correlation between downregulated genes and the X chromosome is dosage compensation. In *Caenorhabditis* nematodes, hermaphrodites/females have two copies of the X chromosome, each of which are partially downregulated to match the dosage of the single X chromosome in males [22]. To determine if whole-scale changes in dosage compensation are responsible for the downregulated pattern in evolved females, we correlated differential gene expression with gene expression levels for dosage compensated and non-dosage compensated X-linked genes [23]. Only 31 (n_male_ = 2, n_female_ = 20, n_shared_ = 9) of the significant genes located on the X chromosome were present in both data sets (Fig. 4). Of these, seven were dosage compensated in wildtype hermaphrodites/females (XX individuals) and three were dosage compensated in both sexes. These genes were further downregulated in evolved individuals, particularly in females, potentially suggesting an expression level lower than males. Nineteen genes in total were not dosage compensated in wildtype hermaphrodites/females, but were downregulated in evolved females by on average 1.4-fold. Similarly, the mean fold decrease in wildtype expression in males (XO individuals) for these same genes is 1.9-fold. Eight genes in total were not dosage compensated in wildtype males and were downregulated in evolved males by an average 2.4-fold. While not definitive, these results point to a potential breakdown in proper dosage compensation accompanying widespread transcriptome downregulation.

**Figure 4.**
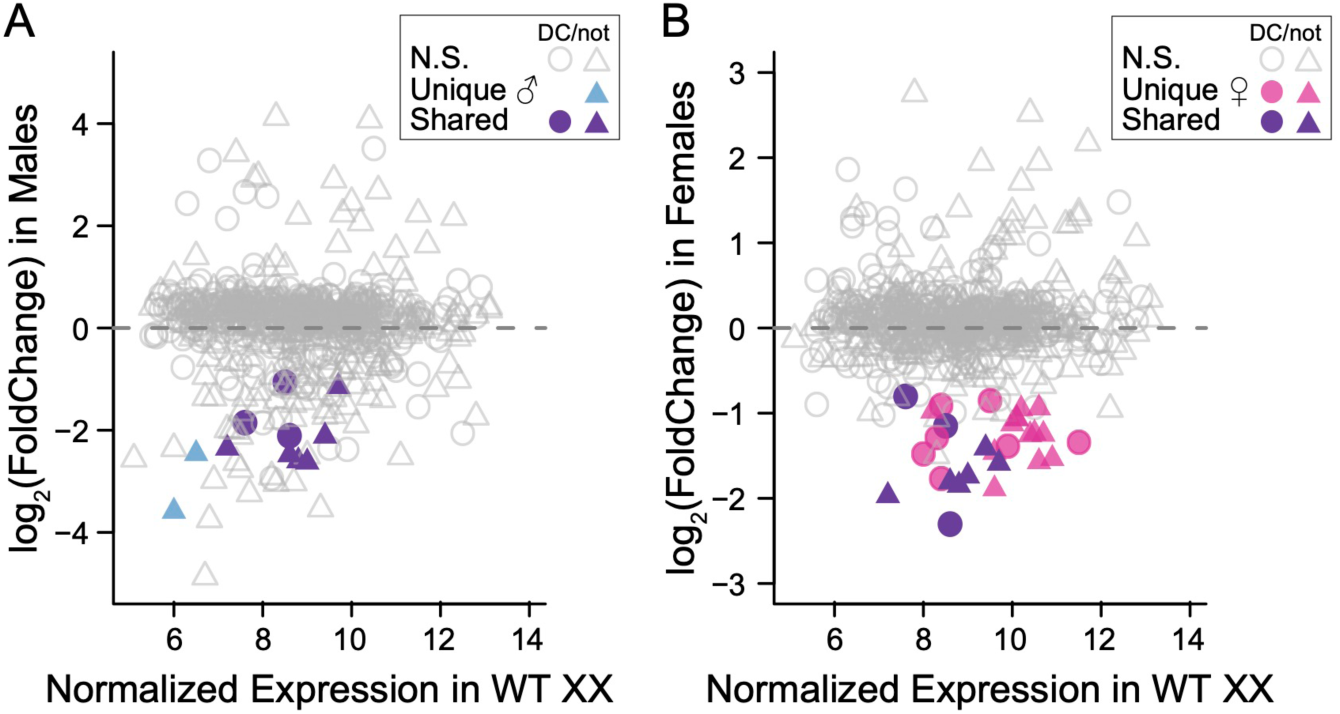
Downregulation of X-linked genes is impacting dosage compensation. **A)** The relationship between normalized expression of dosage compensated (circles) and not dosage compensated (triangles) X-linked genes in wildtype hermaphrodites/females (XX individuals) versus differential gene expression in males. Genes with significant differential gene expression in males only are shown in blue and shared genes are shown in purple. Both axes are plotted on a log_2_ scale. **B)** The relationship between normalized expression of dosage compensated (circles) and not dosage compensated (triangles) X-linked genes in wildtype hermaphrodites/females (XX individuals) versus differential gene expression in females. Genes with significant differential gene expression in females only are shown in pink and shared genes are shown in purple.

The core dosage compensation genes *sdc-2*, *sdc-3*, *dpy-27*, and *dpy-30* did not have significant changes in gene expression after experimental evolution (File S1) nor were there significant allele frequency changes at or near these genes [20]. Additionally, there were no significant allele frequency changes at the *rex* or *dox* binding sites, indicating that the dosage compensation machinery itself is likely not the cause of widespread downregulation in evolved females. Therefore, alternative pathways must be involved in this de-feminization of the male and female transcriptomes resulting from direct sexual selection acting on males.

## DISCUSSION

Sexual selection acts in a sex-specific manner yet can have an indirect evolutionary response in the other sex due to genetic associations created by their sharing the same genome (apart from heterogametic sex chromosomes, when present). When sexual selection acts on one sex, the other sex can be pulled away from their sex-specific optimum as a correlated response to selection, generating conflict between the sexes. Here we explored how the transcriptome of newly evolved males responded to direct sexual selection on sperm competitive ability [20] and whether this male-specific selection generated indirect selection on females by quantifying the ancestral and evolved transcriptomes of males and females mated with different experimental evolution generations.

Genes with significant expression changes between ancestral and evolved males were almost exclusively downregulated. Moreover, most of these genes are female-biased in wildtype individuals. These results suggest that the evolved male transcriptome moved away from a female-like state. This transcriptomic response supports our previous fitness data [20] and indicates that over the course of experimental evolution these newly derived males improved their reproductive ability by becoming more like males under obligate male-female (dioecious) mating. This highlights the first, and fairly ancient, time horizon for sexual conflict revealed by our data. The transition from the ancestral dioecious mating system within *C. elegans* has led to patterns of gene expression that favor hermaphroditic function—which is essentially very female-like—over male-specific function. Selection for increased male function generated in the populations analyzed here [20], as well as that observed in other studies [24–26], leads to a rapid improvement in male reproductive traits such as sperm size and competitive ability. Our results illustrate that these macro-phenotypic effects are reflected directly in the transcriptome via a shift in the balance of expression-level conflict away from female-biased function.

Further, as a byproduct of selection for improved male fitness, females displayed widespread concomitant signature of downregulation in the evolved female transcriptome. This represents sexual conflict on the second time horizon. Selection on regulatory elements that allow enhanced male performance leads to a short-term correlated response to selection on the female transcriptome as well, making it more male-like. Whether we might expect sex-specific transcriptional regulation to rebalance itself over time depends on the specific fitness impacts of these transcriptional changes on females, which are difficult to assess here. Overall, the results indicate a direct de-feminization of the male transcriptome and an indirect masculinization of the female transcriptome generated by direct selection on male fertilization success. Since gene expression levels impact trait function and, in turn, fitness, short-term sexual conflict was likely generated as a byproduct of direct sexual selection in males as female expression was pushed off its global optimum. This give and take is likely to be an important feature structuring transcriptional regulation in many male-female species.

The factorial female mating design used here allows for changes in gene expression to be categorized as a plastic response to increased sperm competitiveness in males or an evolved response in the females themselves. If females were predisposed to increased sperm interactions, then male generation should have been a major contributor to gene expression changes. Such a scenario is analogous to a pre-existing bias model of sexual selection [27] and can lead to direct selection on female preference/resistance [14,15]. However, we found no evidence that male evolutionary history impacted differential gene expression. Rather, all expression changes were determined based on whether a female was from the ancestral or evolved populations, indicating that differential gene expression was a result of indirect selection on females during experimental evolution. Under this scenario optimal gene expression differs between the sexes, such that direct selection optimizing the male transcriptome generates an indirect response in females due to a between-sex genetic covariance in expression [6,13,28]. For our post-insemination model, selection was directed on sperm competitive ability. Since selection on males generated a polygenic response and the expression changes observed were likely due to *trans*-acting pathways, it is possible that multiple reproductive trait may be impacted.

Interestingly, there was a nearly complete lack of overlap between the genic changes identified in Kasimatis et al. (2022) [20] and the expression changes characterized here. One potential reason for this discrepancy is that the genic changes were altering expression of upstream transcription factors that are expressed earlier in development. Kasimatis et al. (2022) [20] identified genomic changes in the coding sequence of or nearby intergenic region of four transcription factors. Of particular note is the transcription factor *sea-1*, which is involved in sex determination and dosage compensation through hypoactivation of the X chromosome [29]. Genomic analysis identified seven SNPs with a significant allele frequency change located within a 30 base pair region of intron 2 [20]. It is possible that this region changes regulation of *sea-1* itself or alters splicing in a way that increase the hypoactivation function of *sea-1* and thus is responsible for the widespread downregulation of X-linked genes. The majority of downregulated genes in females were located on the X chromosome.

Because males are the heterogametic sex in *C. elegans*, alleles affecting gene expression on the X chromosome are directly exposed to selection without any potential shielding of an alternative allele due to dominance, as would occur in heterozygous females. Thus, selection can be more effective on X-bearing genes, especially when there is sex-specific selection [30–32]. The response we observe here is thus consistent with the dominance theory of the rapid evolution of sex chromosomes and their role in speciation [30,33].

Despite the preponderance of signal on the X chromosome, we found no evidence that mutations in the core dosage compensation pathway were the cause of this widespread downregulation. Downregulation of both dosage compensated and non-dosage compensated genes suggests a potential dysregulation of dosage compensation. These changes further support a less wildtype hermaphrodite/female-like state. Wildtype dosage compensation levels may not represent an expression level that is optimal for both sexes [34–36]. Sexual conflict over dosage compensation creates an obvious additional, but perhaps underappreciated, layer to understanding the evolution of sexual dimorphism across the genotype-phenotype-fitness landscape.

Previous experimental evolution studies suggest that male-limited evolution over tens of generations can decrease female fitness [37–39]. Conversely, female-limited or reduced sexual selection evolution tends to increase female fitness with little to no effect on male fitness [40–42]. These previous studies therefore suggest that sex-specific selection acting through males may have a higher potential to generate sexual conflict. However, this previous work has been blind to the traits under selection and therefore the genetic architecture underlying the sexual conflict response. Our work to bridge these layers of the genomic, transcriptomic, and fitness responses suggests a Red Queen scenario [43,44] relating genome evolution and transcriptional dimorphism during post-insemination interactions. Specifically, on microevolutionary timescales strong sexual selection on males can rapidly improve “maleness” at the genomic and phenotypic levels [20] as well as the transcriptomic level, as shown here. In turn, this selection on males pushes females away from their fitness optimum through indirect selection [45,46]. Females may then respond through direct selection, which creates a macroevolutionary picture of a stable female-like state. Thus, examining the functional layers through which sexual dimorphism arises is critical for understanding the context in which sexual conflict can occur.

Sexual selection drives the evolution of some of the most elaborate phenotypic variation and complex behaviors. We show that sexual selection is just as powerful an evolutionary force on post-insemination traits. In particular, on microevolutionary timescales, females may not have the opportunity to counter the effects of indirect selection leading to females being pulled away from their optimal state. Given the seemingly high degree of pleiotropy between female and male reproductive networks, sexual conflict may evolve very rapidly in populations experiencing strong sexual selection.

## MATERIALS AND METHODS

### Worm culture and mating design

We previously evolved six replicated populations of *C. elegans* pseudo-females (*fog-2*) and males under enhanced post-insemination competition for 30 generations [20]. The complete experimental evolution details for the between-strain post-insemination selection only (BS-PO) regime can be found in Kasimatis et al. (2022) [20]. Briefly, evolving males mated with females for 24 hours after which male sterility was induced to prevent further sperm transfer. Fully fertile competitor males were then added to the population to generation sperm competition. After a 24-hour sperm competition period, progeny coming from the evolving males only were collected and propagated to the next generation. Hence, selection was acting on sperm defensive capability and sperm longevity. Here we chose four evolved experimental evolution replicates (A2, A3, B2, B3), which spanned the range of reproductive increase, for transcriptomic analysis.

To generate mated male transcriptomes, we crossed ancestral (strain PX632) females to ancestral (strain PX632) males and evolved (G31 replicates A2, A3, B2, B3) females to evolved (G31 replicates A2, A3, B2, B3) males. The evolved crosses were performed for each experimental evolution replicate. To start a generation, age synchronized larval stage 1 (L1) worms were plated onto a 10 cm NGM-agar plates seeded with OP50 *Escherichia coli* at 20°C with a density of 1,000 worms per plate [47,48]. Late larval stage 4 (L4) females (n = 40) and males (n = 40) were isolated onto medium NGM-agar plates (60 mm diameter) seeded with 100μL OP50 *E. coli*. Three mating plates were set up for each cross. Mating plates were kept at 20°C for 48 hours. After the 48-hour mating period, males were collected and pooled (n = 120) from all three mating plates per cross for bulk RNA isolation. Three biological replicate mating assays were conducted for each cross, each coming from an independent age synchronization event (n = 15 total male samples).

To generate mated female transcriptomes, we used a factorial design of ancestral (strain PX632) and evolved (G31 replicates A2, A3, B2, B3) females mated to ancestral (strain PX632) and evolved (G31 replicates A2, A3, B2, B3) males. Crosses were performed for each experimental evolution replicate following the same experimental procedure as males. After the 48-hour mating period, females were collected and pooled (n = 120) from all three mating plates per cross for bulk RNA isolation. Three biological replicate mating assays were conducted for each cross, each coming from an independent age synchronization event (n = 39 total female samples).

### Transcriptome sequencing, mapping, and gene calling

We performed bulk mRNA sequencing of mated males and females, separately, on day 3 of adulthood. Worms were isolated into 150uL of S-Basal buffer and then pelleted to ∼20uL. They were preserved with 250uL Tri-reagent and flash frozen in liquid nitrogen. The samples went through 10 freeze-thaw cycle before RNA was isolated. RNA was isolated using the KAPA mRNA HyperPrep Kit (KK8580) with Illumina compatible adapters. Libraries were prepared using the TrueSeq RNA Library Prep kit (Illumina) starting from 100 ng of RNA. 100bp single reads for males and 100bp paired-end reads were sequenced on an Illumina HiSeq 4000 at the University of Oregon Genomics and Cell Characterization Core Facility (Eugene, OR).

Reads were trimmed using skewer v0.2.2 [49] to remove low quality bases and TrueSeq adapters (parameters: -x AGATCGGAAGAGCACACGTCTGAACTCCAGTCA-y AGATCGGAAGAGCGTCGTGTAGGGAAAGAGTGT -t 12 -l 30 -r 0.01 -d 0.01 -q 10). The trimmed reads were mapped to the *C. elegans* N2 reference genome (PRJNA13758-WS274) [29] through two passes using STAR v2.5.3a [50]. A table of gene counts was made using HTSeq v0.9.1 [51] (parameters: -s no -r pos -i ID -t gene -f) based on *C. elegans* N2 reference genome “longest isoform” annotations.

### Differential gene expression analysis

To compare-and-contrast changes in gene expression over time based on the generation (*i.e.,* G0 or G31) of individuals at the time of mating, we tested for differential gene expression using the package DESeq2 v1.34.0 [52] in R [53]. DESeq2 uses the generalized linear model framework to estimate the degree to which genes are differentially expressed. We first analyzed each experimental evolution replicate independently. For males, we fit the model: gene counts ∼ male_Anc,Evo_. Since we had a full factorial design for mated females, we fit the model: gene counts ∼ female_Anc,Evo_ + male_Anc,Evo_. In females, experimental evolution replicate B3 had two outlier mating replicates (20_S36_L005 and 21_S37_L005), which were subsequently censored from the dataset (Fig S5). We then combined all experimental evolution replicates to capture only the expression changes that were repeatable across experimental evolution and reanalyzed the differential gene expression model. Normalized count data and gene expression fold change values were extracted from the DESeq2 linear models. However, significance values were determined by fitting a generalized linear model in R with generation and replicate as fixed effects. This approach is more robust to variation between experimental evolution replicates and therefore provides a more conservative estimate significantly differentially expressed genes. Significance was based on a Bonferroni cut-off. These censored, combined data were used for subsequent analyses (Files S1).

We crossed referenced the genes with significant differential expression with the genomic significance peaks identified in Kasimatis et al. (2022) [20]. Additionally, we examined whether the *cis*-regulatory regions of significant differentially expressed genes – defined as 2kb upstream of the start codon – had any SNPs with a significant change in allele frequency, but none of these met the significance peak threshold defined in Kasimatis et al. (2022) [20] (File S2).

We tested for enrichment of gene ontology terms and KEGG (Kyoto Encyclopedia of Genes and Genomes) pathways in significantly differentially expressed genes using the package goseq [54] in R [53]. goseq performs gene ontology analysis taking transcript length biases into account. Median transcript lengths were calculated using from TxDb.Celegans.UCSC.ce11.ensGene [55] based on the UCSC Genome Browser *C. elegans* genome version “ce11”. Gene ontology enrichment was calculated using the “Wallenius” method with a 0.05 false-discovery rate cutoff [56] (File S3).

### Sex-biased gene expression and dosage compensation

We compared the change in expression of significant genes with their wildtype expression profiles in *C. elegans fog-2* females and males [21] to examine any potential changes in sex-biased expression profiles. The relationship between sex-biased expression and differentiation gene expression in each sex was analyzed by fitting a linear model in R.

The dosage compensation profiles for *C. elegans* hermaphrodites (XX individuals) and males (XO individuals) were taken from Jans et al. (2009), as were the positions of *rex* and *dox* binding sites. We then correlated dosage compensation profiles with all genes in our transcriptomic dataset. The positions of *rex* and *dox* sites were compared with genomic SNP data from Kasimatis et al. (2022) [20] to determine if any significant allele frequency changes occurred in these regions.

## Data Accessibility

Raw transcriptomic reads are available at NCBI SRA under accession number PRJNA1013082. Differential expression analysis results for the combined data are in File S1. Cross-referencing of *cis*-regulatory SNPs are available in File S2. Gene ontology and KEGG pathway enrichment results are available in File S3. All scripts are publicly available on the GitHub repository: https://github.com/katjakasimatis/expevol_femaletranscriptomics. The experimental evolution ancestral strain (PX632) and evolved replicates are available from the Phillips lab upon request.

## Acknowledgements

We thank Ruben Lancaster and Alex Smith for experimental assistance and Anastasia Teterina for transcriptome analysis advice. We thank Locke Rowe and Stephen Wright for their insightful discussion.

## Author Contributions

KRK and PCP devised the project. KRK and CS performed the crosses. JHW prepared the transcriptomic libraries. KRK analyzed the data. KRK wrote the manuscript with the support of the other authors.

---

**Figure S1.**
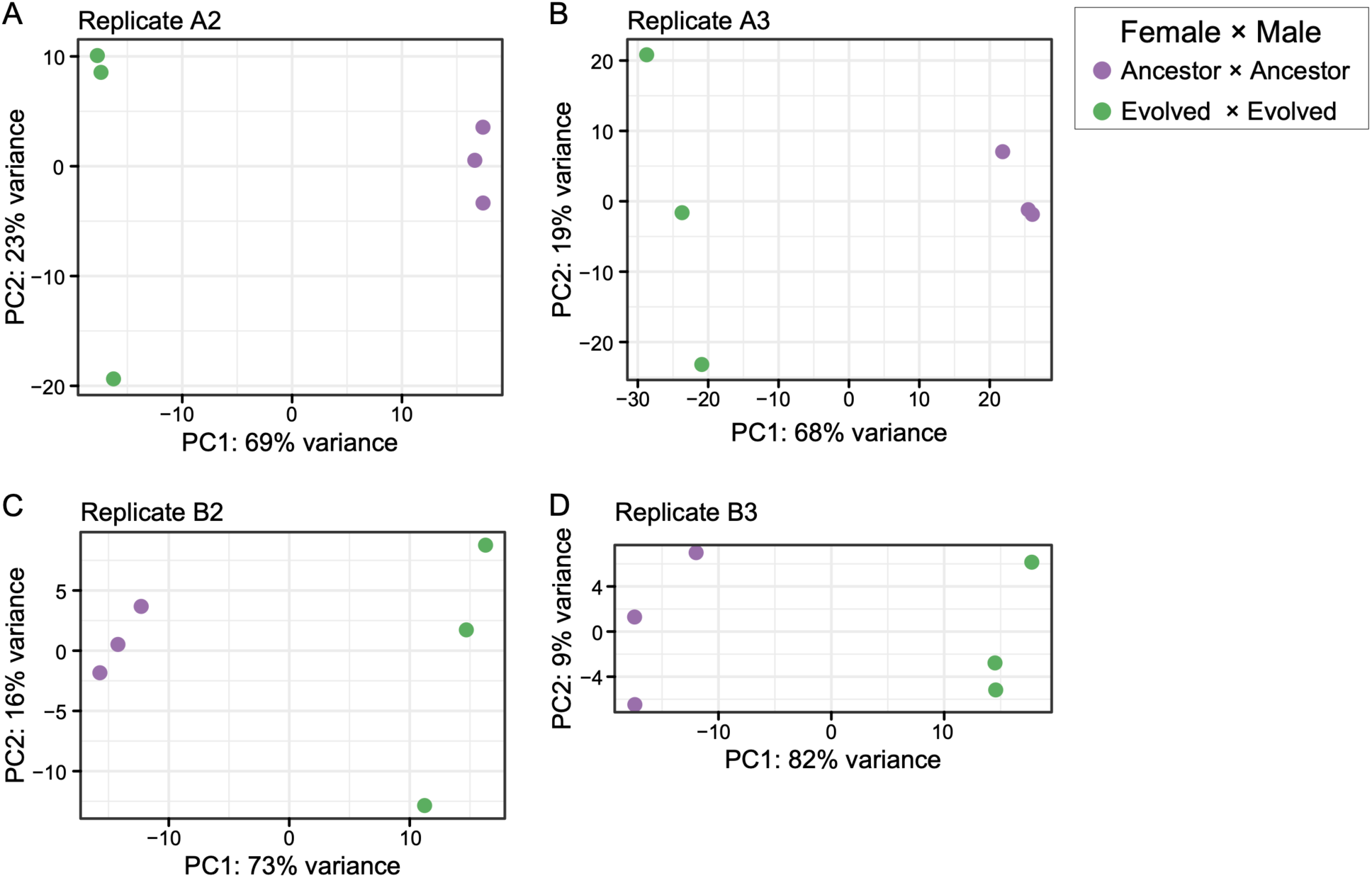
Principal component analysis of the variance stabilized data for each experimental evolution replicate partitions male generation along PC1.

**Figure S2.**
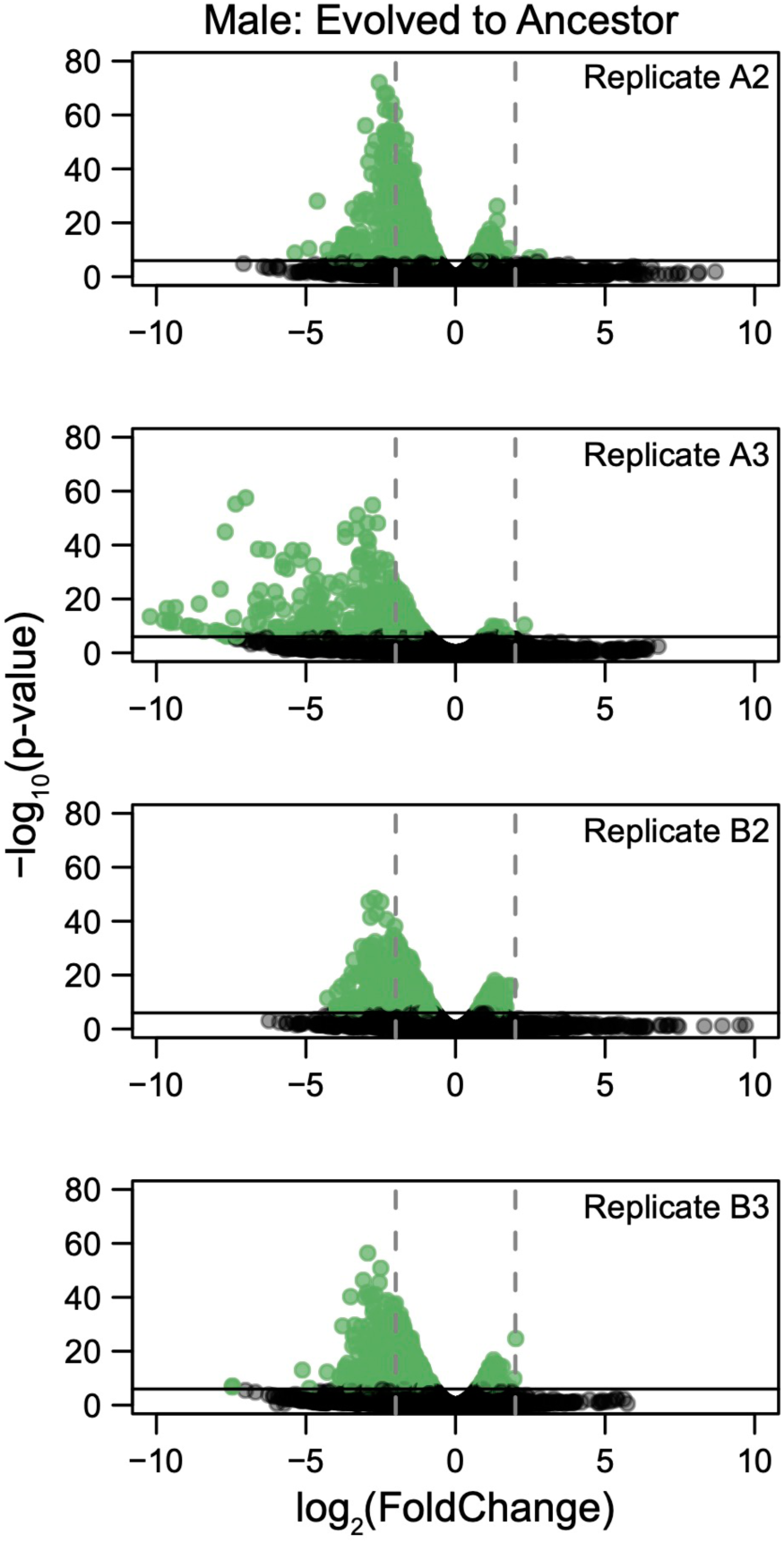
Differential gene expression in males for each experimental evolution replicate was analyzed based on male generation. Genes with a significant change in expression from ancestral to evolved males (shown in green) were determined using a Bonferroni cut-off. Positive expression changes correspond to upregulation in evolved males, while negative expression changes correspond to downregulation in evolved males.

**Figure S3.**
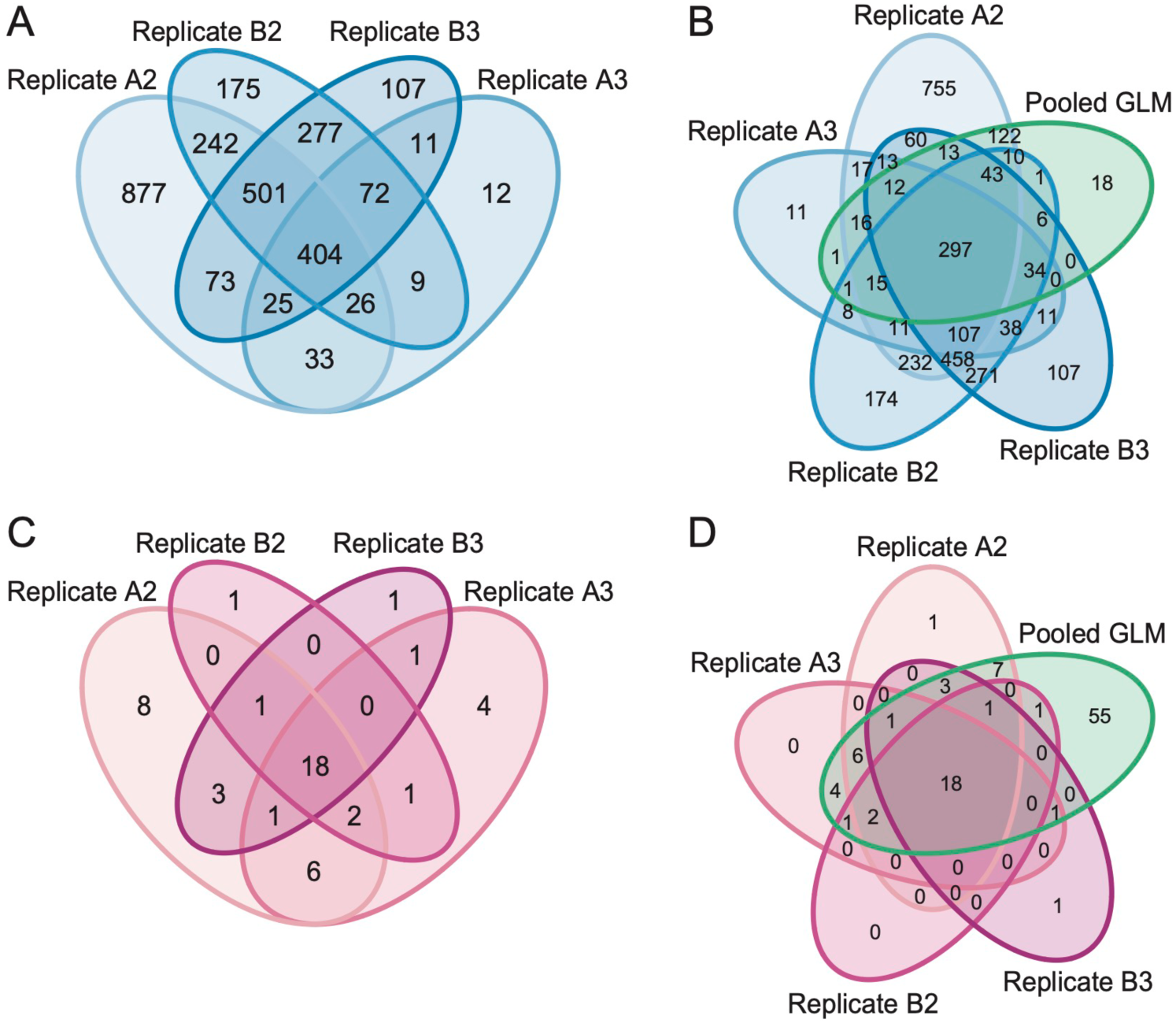
The overlap of significant genes across individual experimental evolution replicates in: **A)** males and **C)** females. The overlap between the individual experimental evolution replicate analyses and the combined analysis in: **B)** males and **D)** females.

**Figure S4.**
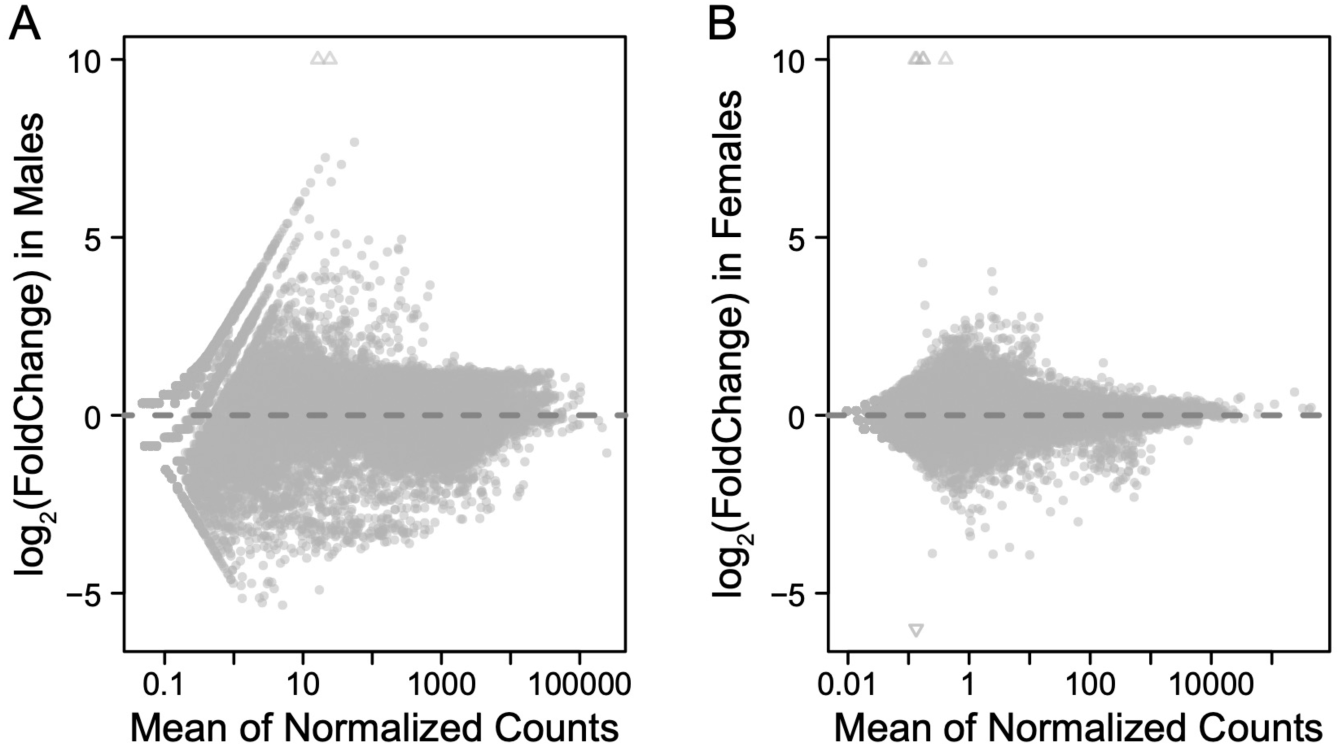
Distribution of normalized gene expression counts against the change in gene expression for the combined analysis of: **A)** males and **B)** females. Triangles represent points that fall outside of the y-axis range.

**Figure S5.**
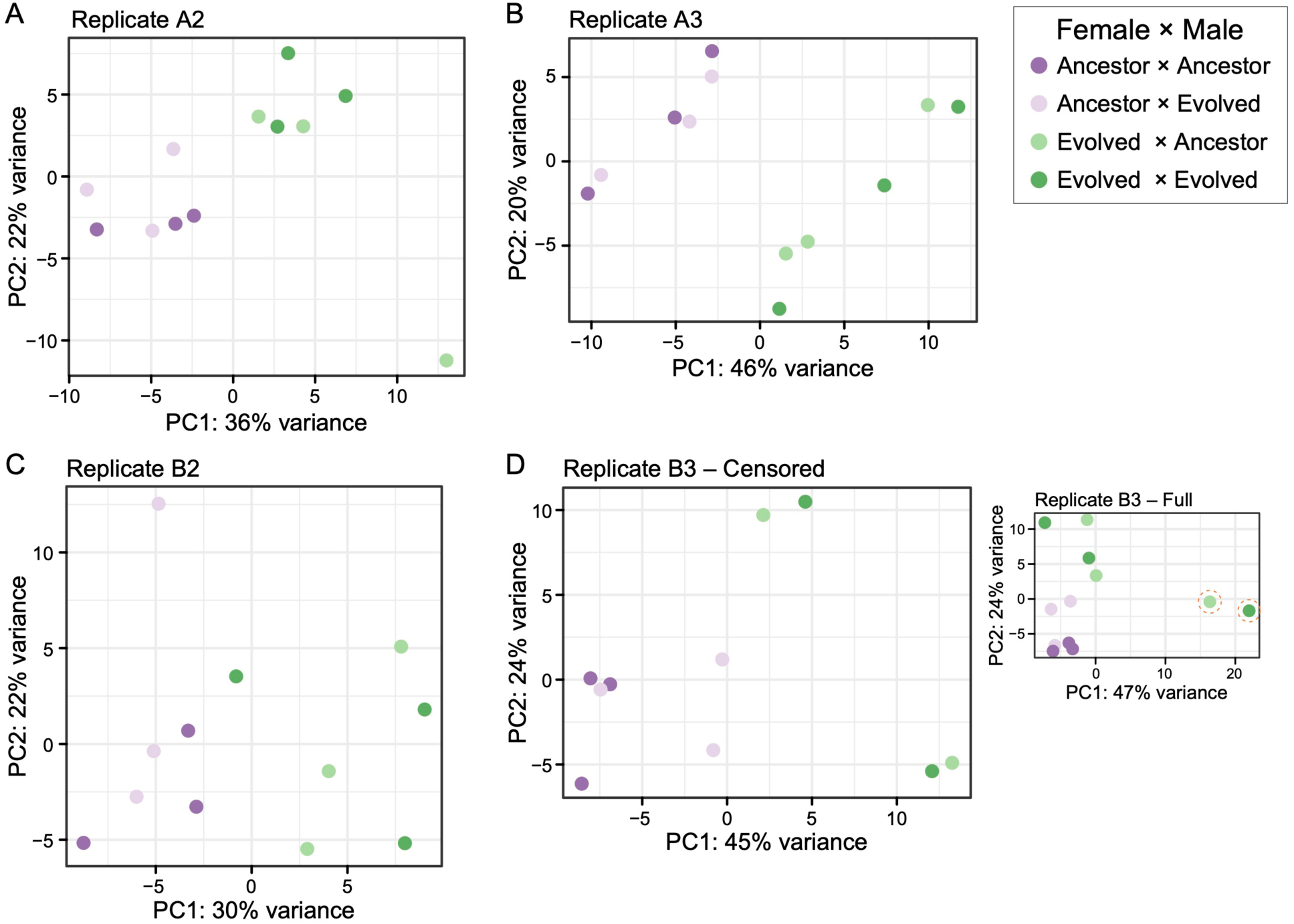
Principal component analysis of the variance stabilized data for each experimental evolution replicate partitions female generation along PC1. Experimental evolution replicate B3 had two outlier transcriptome replicates (circled) and were censored from the final, combined dataset.

**Figure S6.**
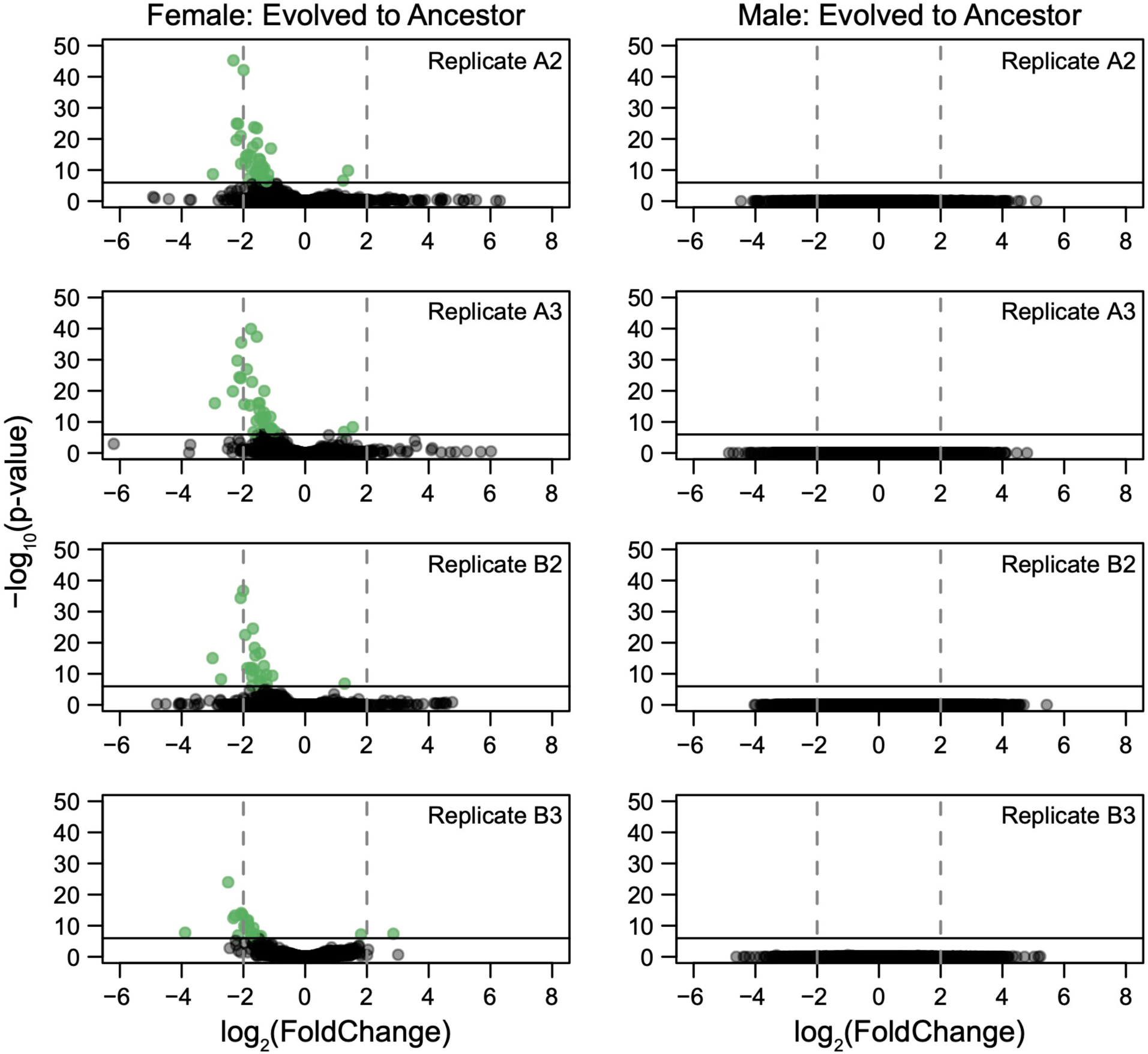
Differential gene expression in females for each experimental evolution replicate was analyzed based on female generation and male generation. Genes with a significant change in expression from ancestral to evolved females (shown in green) were determined using a Bonferroni cut-off. Positive expression changes correspond to upregulation in evolved females, while negative expression changes correspond to downregulation in evolved females.

**Figure S7.**
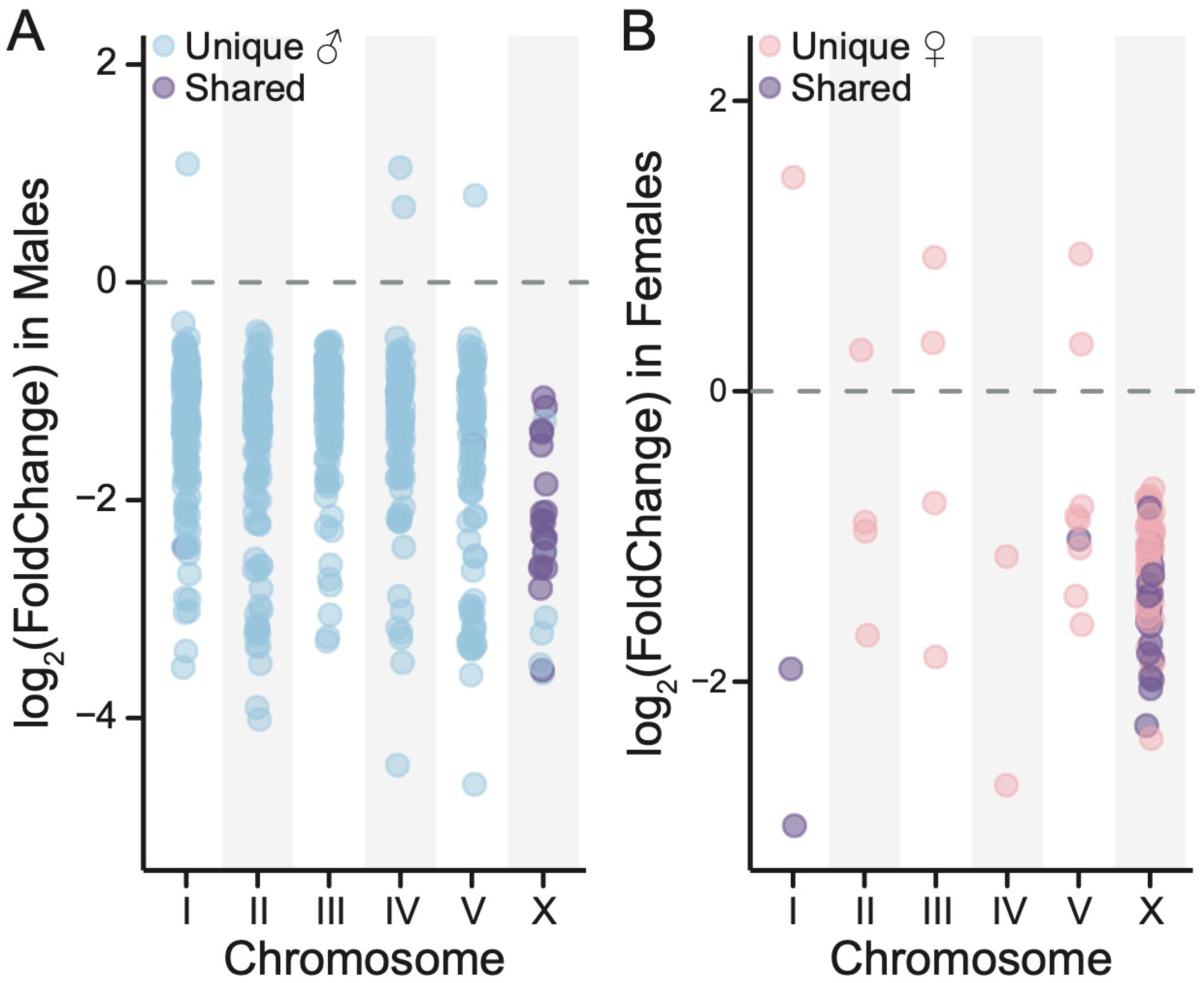
Chromosomal location of significantly differentially expressed genes. **A)** In males, the majority of DEs are located on the autosomes. Unique male DEs are shown in blue and shared DEs are shown in purple. **B)** In females, the majority of DEs are located on the X chromosome. Unique female DEs are shown in pink and shared DEs are shown in purple.

**File S1.** Differential expression analysis for the combined male and combined female datasets.

**File S2.** Genomic SNPs located in the cis-regulatory region of DEs.

**File S3.** Gene ontology analysis and KEGG Pathway analysis for DEs.

